# Distinguish virulent and temperate phage-derived sequences in metavirome data with a deep learning approach

**DOI:** 10.1101/2020.12.25.424404

**Authors:** Shufang Wu, Zhencheng Fang, Jie Tan, Mo Li, Chunhui Wang, Qian Guo, Congmin Xu, Xiaoqing Jiang, Huaiqiu Zhu

## Abstract

**Background:** Prokaryotic viruses referred to as phages can be divided into virulent and temperate phages. Distinguishing virulent and temperate phage-derived sequences in metavirome data is important for their role in interactions with bacterial hosts and regulations of microbial communities. However there is no experimental or computational approach to classify sequences of these two in culture-independent metavirome effectively, we present a new computational method DeePhage, which can directly and rapidly judge each read or contig as a virulent or temperate phage-derived fragment.

**Findings:** DeePhage utilizes a “one-hot” encoding form to have an overall and detailed representation of DNA sequences. Sequence signatures are detected via a deep learning algorithm, namely a convolutional neural network to extract valuable local features. DeePhage makes better performance than the most related method PHACTS. The accuracy of DeePhage on five-fold validation reach as high as 88%, nearly 30% higher than PHACTS. Evaluation on real metavirome shows DeePhage annotated 54.4% of reliable contigs while PHACTS annotated 44.5%. While running on the same machine, DeePhage reduces computational time than PHACTS by 810 times. Besides, we proposed a new strategy to explore phage transformations in the microbial community by direct detection of the temperate viral fragments from metagenome and metavirome. The detectable transformation of temperate phages provided us a new insight into the potential treatment for human disease.

**Conclusions:** DeePhage is the first tool that can rapidly and efficiently identify two kinds of phage fragments especially for metagenomics analysis with satisfactory performance. DeePhage is freely available via http://cqb.pku.edu.cn/ZhuLab/DeePhage or https://github.com/shufangwu/DeePhage.

## INTRODUCTION

In a microbial community, phages are the major component of the viral genetic materials. It is estimated that the number of phages is on average ten times higher than that of bacteria. They may destroy bacteria, meanwhile in some situations benefit populations of bacteria, and thus crucially impact the microbial community [1]. With the development of the high-throughput sequencing technology, a large number of novel phages are discovered from untargeted metagenomes and viromes, in which viral particles are first enriched before sequencing [2,3]. However, the analysis of these phage sequences is a great challenge since the reference genomes of phages are very limited in view of the fact that most of the phages cannot be cultured independently. The complete phage genomes in current databases are much less than that of bacteria, therefore a large number of sequences from virome data cannot find regions with homology to the known phages [2]. In addition, unlike bacteria, phages lack the universal marker genes such as 16S rRNA [4], so that many species identification strategies designed for bacterial analysis are not applicable to phages. Moreover, for mobile elements such as phages, the sequence assembly is often poorer than that of bacteria, usually because the mobile elements carry repetitive regions like insertion sequences and share sequences that occurred among different genomes [5]. As a result, the large number of short fragments in metagenomic data also increases the difficulty of the analysis.

To overcome these difficulties, several computational tools focusing on two major tasks have been developed to analyze the phage sequences from metagenome or virome. One of the tasks is to identify phage fragments in untargeted metagenomic data, such as the tools VirSorter [6], VirFinder [7], MARVEL [8], virMine [9], and PPR-Meta [10]. Especially, PPR-Meta is a tool with high performances developed by us and demonstrates much better accuracy than the related tools.

Another task is to assign the host for a given phage contig, such as the tools WIsH [11], VirHostMatcher [12], and Hostphinder [13]. However, these tools cannot answer the question about how the discovered phages interact with their hosts. According to the interaction mode, which is also referred to as the phage lifestyle, phages can be divided into the virulent phages and the temperate phages [14]. When a virulent phage infects its host, it will produce many progenies as soon as the phage’s DNA is injected into the host cell and then cause the death of the host through bacterial lysis [14]. In contrast, temperate phages can undergo the lysogenic cycle and lytic cycle. In the lysogenic cycle, a temperate phage will integrate its genome into the host chromosome, which is also referred to as prophage, and then copies its genome together with the host chromosome [15]. While induced by appropriate conditions, especially the nutritional conditions and the number of co-infecting phages, temperate phages will go into the lytic cycle, following by releasing the viral particle and killing the host through bacterial lysis [16]. Such different processes have a significant influence on the microbiota especially in the human gut, which could be highly correlated with human diseases or the treatment of human disease. Although some kinds of hotspots, such as phage therapies that making use of the virulent phages in the context of therapeutic use [17], have been investigated, limited by current bioinformatics tools, people still knew little about these different lifecycles for their prevalence in the human gut [18]. Therefore, it is important to distinguish virulent and temperate phages for further understanding of phage-host interaction.

Although the classification strategy of this issue for virome data has not been investigated yet, there are several noteworthy works that help to characterize the virulent and temperate phages. Even phages lacking marker genes, those studies show that they may have some functional genes, which are high-frequency genes and can tell us whether a given phage is virulent or temperate in a relatively credible way. For example, Emerson et al. found there were some functional genes for temperate phages, such as integrase and excisionase [19]; Schmidt et al. found that the leucine substitution in DNA *polA* gene had a strong connection with temperate phages [20]. Notably, McNair et al. designed a computational tool called PHACTS to identify whether a phage with a complete or partial proteome is virulent or temperate [14]. This tool employs all the sequence information of proteins from a phage genome and uses the random forest as a classifier to make the judgment. Researchers further found that the existence of some kind of genes helped PHACTS present good results. For example, virulent phages usually have genes related to phage lysis, nucleotide metabolism, or structural proteins, while temperate phages usually contain genes related to toxins, excision, integration, lysogeny, or regulation of expression [14]. Unfortunately, such kind of strategies may not apply to metagenomic data. To date, it is still a difficult task to reconstruct complete genomes of all organisms in the metagenomic data. Therefore, only a few DNA fragments may contain those functional genes that can help to make the judgment. According to the report of PHACTS, this tool can achieve accuracy over 95% if at least 25 proteins are provided from a phage genome. However, if fewer proteins are obtained, the accuracy of PHACTS reduces obviously. When only five proteins from a phage, PHACTS only achieves an accuracy of about 65%; if only two proteins from a phage, PHACTS appears to produce random results with an accuracy below 55%. Considering that most of the DNA sequences in metagenomic data are short fragments that only contain a few genes or even incomplete genes, it is essential to develop a tool, which does not depend on using information from sufficient proteins with functional genes level, while to make judgment directly for each short DNA fragment in metagenomic data.

In this paper, we present a two-class classifier DeePhage to identify whether a DNA fragment is derived from a virulent phage or a temperate phage. Using the information of every nucleotide without manually feature extracted, DeePhage encodes sequences in “one-hot” form. Such representations are suitable for the Convolutional Neural Network (CNN) model to detect helpful motifs for classification, which are common used on biological sequence identification. Together with other kinds of neural network layers, DeePhage learns different features between virulent and temperate sequences and then outputs a score indicating the possibility to be a certain kind of phage sequence. Tested on the same data, DeePhage can significantly outperform the best of available methods PHACTS on computational efficiency by using only 1/810 computation time that PHACTS uses. Simulation studies on five-fold validation show that DeePhage precedes by approximately 30% compared with PHACTS. DeePhage’s evaluations on real metavirome data of bovine rumen are better than PHACTS with much more accurate results, which use annotations of the BLAST method as a reference. Meanwhile, we present a new strategy to conveniently detect the phage transformation by tracing specific phage contigs, which can explore the influence of phages that contribute to human diseases. DeePhage can be used to analyse the virome data and untargeted metagenomic data directly. While handling the metagenomic data, users need to firstly identify the phage sequences using related software, such as PPR-Meta [10] as we mentioned above, and then use DeePhage to further annotate the phage sequences.

## MATERIAL AND METHODS

### Data construction

Considering that there is no real virome data with the reliable lifestyle annotation for each sequence as a benchmark, we constructed artificial contigs extracted from well-annotated complete phage genomes as the benchmark to train and test the algorithm. We downloaded 227 complete phage genomes with lifestyle annotations from McNair dataset, including 79 virulent phages and 148 temperate phages [14]. Among these phages, we removed two virulent phages from the dataset: mycobacteriophage D29 (accession: NC_001900) and lactococcus lactis bacteriophage ul36 (accession: NC_004066), because the lifestyle of these two phages may be ambiguous. Although these two phages are annotated as virulent phages, researchers found that they both contained functional integrases, indicating that they can integrate their genomes into host chromosomes like temperate phages [14]. Besides, D29 is very similar to the temperate phage L5 [21], while ul36 has 46.6% homology with the temperate phage Tuc2009 [22]. Therefore, 77 virulent phages and 148 temperate ones are used in the current study. In general, the unbalanced size between positive and negative samples may have an impact on the accuracy of the machine learning-based algorithm [7]. In the McNair dataset used in this work, it is thus obvious that the number of positive samples is less than that of negative samples. However, we found that the genome length of each positive sample is generally longer than that of each negative sample. It is probably because temperate phages can integrate their genomes into host chromosomes and may discard some non-essential genes. What is more, genes on host chromosomes may be served as compensation. As a result, based on the bases counts, the dataset size between positive samples and negative samples are similar. For convenience, herein the virulent phages are referred to as the positive sample and the temperate phages as the negative sample.

We further used MetaSim (v0.9.1) [23] to extract artificial contigs from the complete phage genomes. Considering that the length of contigs in real metagenomes may cover a wide range, we divided the artificial contigs into four groups according to their length: the length range in Group A is 100-400 bp; Group B is 400-800 bp; Group C is 800-1200 bp while Group D is 1200-1800 bp. Those four groups may cover the length of raw reads and the average length of assembled contigs from the next-generation sequencing technology. We would evaluate the performance of DeePhage on different groups respectively.

We also used real virome data to estimate the reliability of DeePhage qualitatively. We downloaded virome data of bodily fluid in the bovine rumen [24] from MG-RAST [25]. They were downloaded as raw reads (accessions: mgm4534202.3 and mgm4534203.3). We used SPAdes (v3.13.0) [26] to assemble the raw reads and obtained 118918 contigs with the N50 of 291 bp.

### Mathematical model of phage sequences

To evaluate the feasibility of using sequence signature used for classifying virulent and temperate phages, we first analysed the distribution of k-mer frequencies, which have been widely used to distinguish genomes from different species, among virulent phage genomes and temperate phage genomes. We used 4-mer frequencies to characterize each phage genome in our dataset. The Principal Component Analysis (PCA) [27] revealed that the 4-mer frequencies between virulent and temperate phage genomes have different distribution (Figure 1), showing that they have different sequence signatures to characterize these two categories of phage genomes.

**Figure 1.**
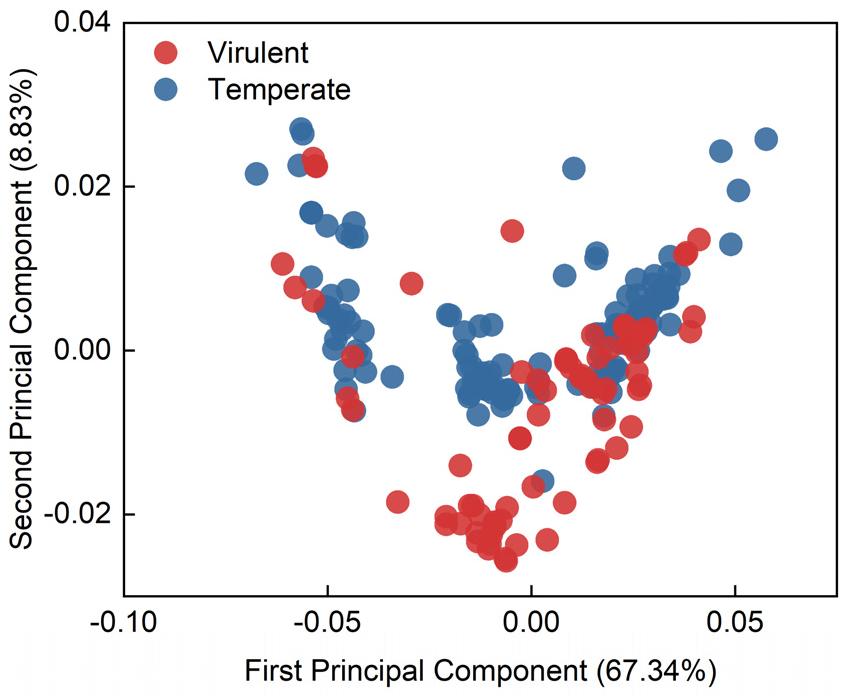
The PCA of 4-mer frequencies distribution among virulent and temperate phage genomes.

Although k-mer frequencies have shown their ability to classify virulent and temperate phage genomes, using such frequencies to characterize short DNA fragments will usually be disturbed with the noise (11). Also, as global statistics that may miss some local information, k-mer frequencies are difficult to detail characterize mobile elements that contain mosaic structure [28]. To describe the local sequence information in detail, we consider the one-hot encoding form, which can represent every base continuously and entirely. For each sequence, we used the “one-hot” encoding form to represent each base in a sequence. Specifically, bases A, C, G and T were represented by [0,0,0,1], [0,0,1,0], [0,1,0,0], and [1,0,0,0].

### Algorithm structure of DeePhage

Deep learning algorithms are recognized as an extremely effective method in many fields including in the biology field. Comparing with the Recurrent Neural Network (RNN), the Convolutional Neural Network (CNN) models are faster to train and more efficient in sequential spatial correlations [29]. Specifically, CNN is a universal network for extracting local patterns in terms of biology, which in the current context can be used as a motif detector of DNA sequences. In DeePhage, we presented a deep learning algorithm with CNN models to handle the input sequences represented by the one-hot encoding form. The network contained eight layers: a 1D convolutional (Conv1D) layer, one 1D maximum pooling (Maxpooling) layer, one 1D global average pooling (Globalpooling) layer, two batch normalization (BN1 and BN2) layer, a dropout (Dropout) layer, and two dense (Dense1 and Dense2) layers.

Conv1D layer takes a sequence encoded by an L×4 matrix *X* (L is the length of the sequences, equals 400, 800, 1200, and 1800) as the input and generates total F feature maps as output by corresponding F convolutional kernels. Using ReLU (Rectified Linear Unit) [30] as the activation function, the Conv1D layer output an L×F matrix *Y^c^* and computes for the f^th^ feature map at the l^th^ location like this:

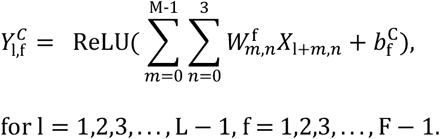

The *W^f^* and 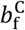 are an M×4 weight matrix and a bias of the f^th^ kernel. The mentioned ReLU function is defined as [30]:

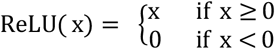

As a traditional nonlinear function, the ReLU function is easier to train and achieves better performance, which can rectify the shortcomings of sigmoid functions. Those kernels scan a sequence one after another to extract the valuable features for the classification and the ReLU function achieves a nonlinear transformation.

Such a combination is followed by the Maxpooling layer to downsample the input representation by taking the maximum value over an input channel with a pooling size S1 and a stride size S2. The window is shifted along with each channel independently and can generate F new channels with the size of L’ (L’ = L/S2). The Maxpooling layer outputs an L’ ×F feature matrix *Y^M^* and one of the pooling operation for a specific channel at the l^th^location defines like this:

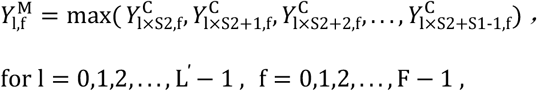

Its main function is to reduce the dimensions of each input channel using final summarised features, which can also adapt to location variations of valuable features.

Features from the Maxpooling layer are passed to the BN1 layer to scale the inputs. At each batch, it usually transforms inputs to have a mean close to 0 and a standard deviation close to 1, which can avoid the vanishing gradient problem and accelerate the convergence rate of the model. So the output feature matrix *Y^B1^* of the BN1 layer is also an L’ ×F matrix as *Y^M^* but being scaled.

The next is a Dropout layer, which randomly drops a certain proportion (denoted as P) of input elements by setting them to zero during training (29). The output ***Y***^Dp^ is formulated as:

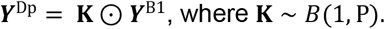

The drop mask **K** denotes a Bernoulli distribution with *n* equals 1 and *p* equals P. It could effectively reduce overfitting especially in our small dataset [31].

After a dropout layer, the Globalpooling layer takes the ***Y***^Dp^ as input and reduce features from the same channel into one dimension by using the average value of those features, which can integrate global spatial information. More formally:

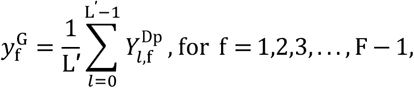

where 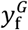 is the average value of features from the f^th^ input channel. Considering all the F channels from the previous layer, the output of the Globalpooling layer ***y**^G^* is an F dimension vector.

Subsequently, a Dense1 layer using ReLU function as activation function outputs R units. It has an R×F weight matrix *W*^D1^ and an R-dimensional bias vector *b*^D1^. Each output units is processed by:

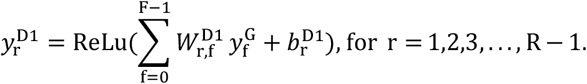

Dense1 layer can compile the features from different input channels together and finally generate an R-dimensional vector *y*^D1^, while a Conv1D layer just extracts features into different feature maps.

The vector *y*^D1^ is then sent into a BN2 layer to generate a new feature vector *y*^B2^ that having a mean close to 0 and a standard deviation close to 1, which has the same effect as the BN1 layer.

Using a sigmoid function as an activation function, the final layer is the Dense2 layer and output only one score between zero and one representing the probability of prediction. Using an R-dimensional weight vector *W*^D2^ and a bias scalar *b*^D2^, the output score is given by:

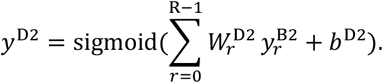

The sigmoid function is defined as:

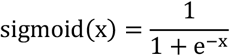

In general, the sequence with a score higher than 0.5 would be regarded as a positive sample (a virulent phage) and the sequence with a score lower than 0.5 would be regarded as a negative sample (a temperate phage). When training, we used the Adam optimizer [32] (learning rate = 0.0001), binary cross-entropy as the loss function, and 32 as the batch size to train the neural network and update network weights. Altogether, we found that setting the size F to 64, M to 6, S1 to 3, S2 to 3, P to 0.3, and R to 64 made the best performance. The structure of DeePhage was shown in the upper part of Figure 2.

**Figure 2.**
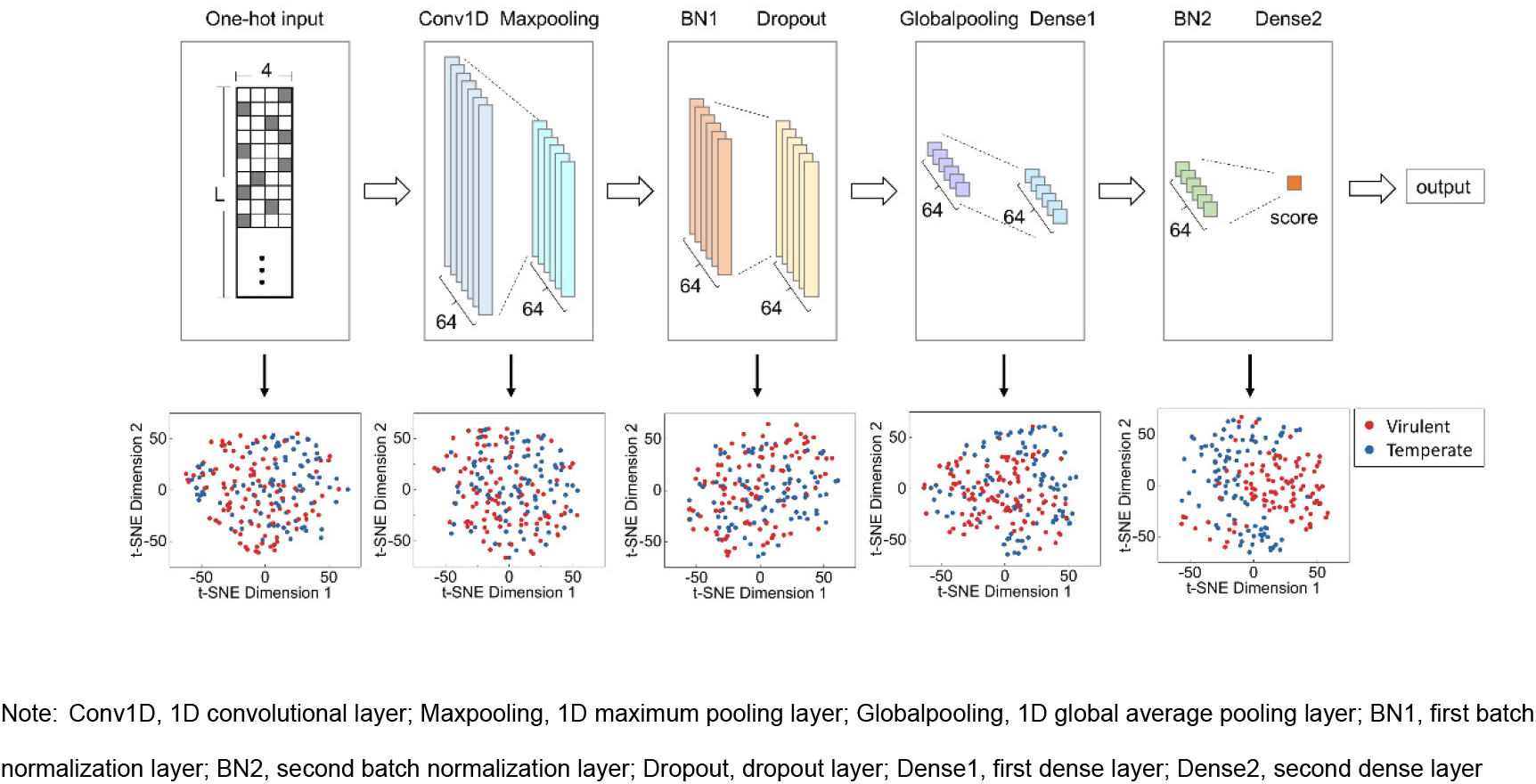
Structure of deep learning neural network and visualization of five layers by reducing dimensions. DeePhage uses the Convolutional Neural Network as the classifier. The neural network (in the upper part) takes the sequence in the “one-hot” coding form as input and output a score between zero and one. In general, the sequence with a score higher than 0.5 can be referred to as the virulent phage-derived fragment and the sequence with a score lower than 0.5 can be referred to as the temperate phage-derived fragment. The visualization demonstrated the learning process of DeePhage. The performance would be better when we observing a deeper layer (in the lower part).

It is worthy to know more about the importance of the encoding method for sequences and each specific layer in our model, so we tested six different models (including DeePhage) by using k-mer frequencies as an encoding representation or removing a certain layer. The six model architectures (DeePhage, Kmer-4, No-Maxpooling, No-Dropout, No-Globalpooling, and No-BN) were shown in Additional File 1 (Figure S1) and their performances were shown in Additional File 1 (Table S1). It can be seen that the Kmer-4 model did a terrible prediction. As mentioned above, when we used 4-mer frequencies to characterize each phage at the level of genome sequences (as shown in Figure 1), it could slightly distinguish two kinds of phages. It is proved that k-mer frequencies have not enough power to represent short sequences and are fit for capturing the global signature of long sequences rather than the local signature of short sequences. As for those models removing a certain layer, the performance dropped compared with DeePhage. Especially, the prediction accuracy reduced nearly 10% and 5% when using a model without a Globalpooling layer and BN layers (No-Globalpooling and No-BN model). Other models decreased slightly. We can see the architecture and the one-hot encoding representation of DeePhage are better than others.

Although deep neural networks are considered as black-box models, we hope to have insights about the learning process for features. We chose five layers (One-hot input, Conv1D, BN1, Globalpooling, and BN2) to observe their learned features. Because it is hard to gain an intuitive feeling about high-dimension features, we used t-Distributed Stochastic Neighbor Embedding (t-SNE) [33], which is a machine learning algorithm for dimensionality reduction, for the visualization of high-dimensional data in a 2D projected space. During the training period, we firstly used PCA to reduce features into a 20-dimensional space and then used t-SNE to reduce them into a two-dimensional space using the sequences from Group D. The visualizations of five layers were shown in the lower part of Figure 2. It could be seen that the effects of classification are better when focusing on the deeper layers. In detail, two types of phages were firstly mixture together and then separated gradually, which demonstrated the learning process of DeePhage. Furthermore, it should be emphasized that the visualizations by dimensionality reduction cannot reflect the complete power of DeePhage.

Considering the other length of sequences beyond our four-trained groups, we design some strategies. For those sequences longer than 1800 bp, DeePhage will split the sequence into several 1800-bp-long subsequences without overlapping, usually except the last subsequence. DeePhage will then use the neural network in the corresponding group to predict each subsequence, and calculate the weighted average score according to the score and length of each subsequence. Because training the neural network using long sequences is very time-consuming, we do not train additional neural networks for longer sequences. For those sequences shorter than 100 bp, DeePhage uses the neural network in Group A to predict.

## RESULTS

### Identification performance of DeePhage

We first used the five-fold cross validation to evaluate the performance of DeePhage. To test whether DeePhage can distinguish the lifestyle of novel phages or not, for each validation, we divided the training set and the test set based on complete genomes rather than artificial contigs, and then simulated 20,000 training sequences and 20,000 test ones using MetaSim [23]. The performance evaluation criteria here are defined as: *Sn=TP/(TP+FN); Sp=TN/(TN+FP)*; and *Acc=(TP+TN)/(TP+ TN+FN+FP)*. Among these criteria, *Sn* and *Sp* are used to evaluate the accuracy of virulent phages and temperate phages respectively, while *Acc* is used to evaluate the overall performance. As shown in Table 1, DeePhage demonstrates overall reliable and stable performance with *Acc* from 75% to 88%. Such results indicate that the input of functional genes with several proteins is not required for our DeePhage. Therefore, our DeePhage method shows an evident advantage compared with the tool PHACTS. Since DeePhage can identify each DNA fragment as the virulent phage-derived sequence or temperate phage-derived one directly and independently, it would be a more acceptable tool to analyse phages in metagenomic data. In this case, the complete or near-complete genomes for phages were hard to be reconstructed from the data, especially for those with low abundance or in a low coverage sequencing condition. Clearly, our DeePhage has the advantage of being applicable to processing the data by current short-read sequencing technologies and performs better when the short reads could be assembled into longer contigs.

**Table 1.** Results of five-fold cross-validation for DeePhage. The validation of each group was performed independently. Each result consists of the mean and standard deviation.

| Group | Group A<br>(100-400 bp) | Group B<br>(400-800 bp) | Group C<br>(800-1200 bp) | Group D<br>(1200-1800 bp) |
| --- | --- | --- | --- | --- |
| *Sn*(%) | 69.3±3.6 | 76.2±5.5 | 82.1±6.6 | 86.9±4.7 |
| *Sp*(%) | 78.8±5.8 | 86.1±6.9 | 88.6±9.6 | 88.7±8.9 |
| *Acc*(%) | 74.6±1.9 | 81.8±2.7 | 85.8±4.0 | 88.2±4.3 |

It is noted that the *Sn* is slightly lower than *Sp* for each rotation of different length groups, which means that some virulent phage-derived sequences are more prone to be misjudged as temperate ones. The reason is possibly that the diversity of virulent phages is lower than that of temperate phages in the current database. Although the sizes of positive and negative samples are comparable based on base counts, the number of genomes in positive samples is less than that of negative samples. In general, sequences from the same genome have similar sequence signatures such as codon usage and GC content. The fewer number of genomes in positive samples may lead to lower diversity. From the algorithm of machine learning, the negative samples have a wider distribution in the feature space while the positive samples only occupy a smaller space. Therefore, for the test data, the positive samples are easier to fall out of their feature space, which leads to the misjudgment of DeePhage. Despite this, the performance of DeePhage on virulent phages is rather reliable. More details about the performances of the ROC curves and AUC scores of DeePhage in each rotation of the five-fold cross validation are shown in Additional File 1 (Figure S2).

In general, sequences with scores near 0.5 are not as reliable as those sequences with a score near 0 or 1. Therefore, DeePhage is designed with an adjustable cutoff to filter out these uncertain predictions. Users can specify a cutoff using a parameter. In this way, a sequence with a score between (0.5-cutoff/2, 0.5+cutoff/2) will be labelled as “uncertain”. In general, with a higher cutoff, the percentage of uncertain predictions will be higher while the remaining predictions will be more reliable.

### Comparison with PHACTS for protein sequence identification

It should be noted that DeePhage and PHACTS were designed for different tasks, PHACTS was designed for complete genomes while DeePhage is designed for metagenomic fragments. Therefore, the requirements of the input data for them are actually different. PHACTS requires users to input all proteins (amino acid sequences) within one phage genome, so proteins from different phages should not be put into the same file. In contrast, the DeePhage’s requirement is only to input all DNA fragments (nucleic acid sequences), no matter whether they contain coding regions and whether they are from the same phage, and DeePhage may directly judge each fragment independently. Although it is difficult to compare two tools based on the same condition, we tried to test the performance of PHACTS in DNA short fragments. Since PHACTS requires a collection of protein sequences as input, we firstly annotated the protein sequences of 100,000 DNA sequences of the test set in each length group using FragGeneScan (v1.31) [34] and proteins from the same sequence (sequences without coding regions were ignored) are input into the program PHACTS (v0.3). As for comparison, DeePhage is also used to predict these DNA sequences with coding regions. The total accuracy (the number of correct predictions divided by the total number of sequences having the coding regions in each length group) of DeePhage and PHACTS in each length group are shown in Figure 3. For short fragments covering data sets of Group A to D, PHACTS demonstrates the accuracies of *ACC* around 50%, which are nearly the results of random predictions. In construct, DeePhage can satisfactorily classify the sequences with the accuracies of *ACC* about 75%~88%.

**Figure 3.**
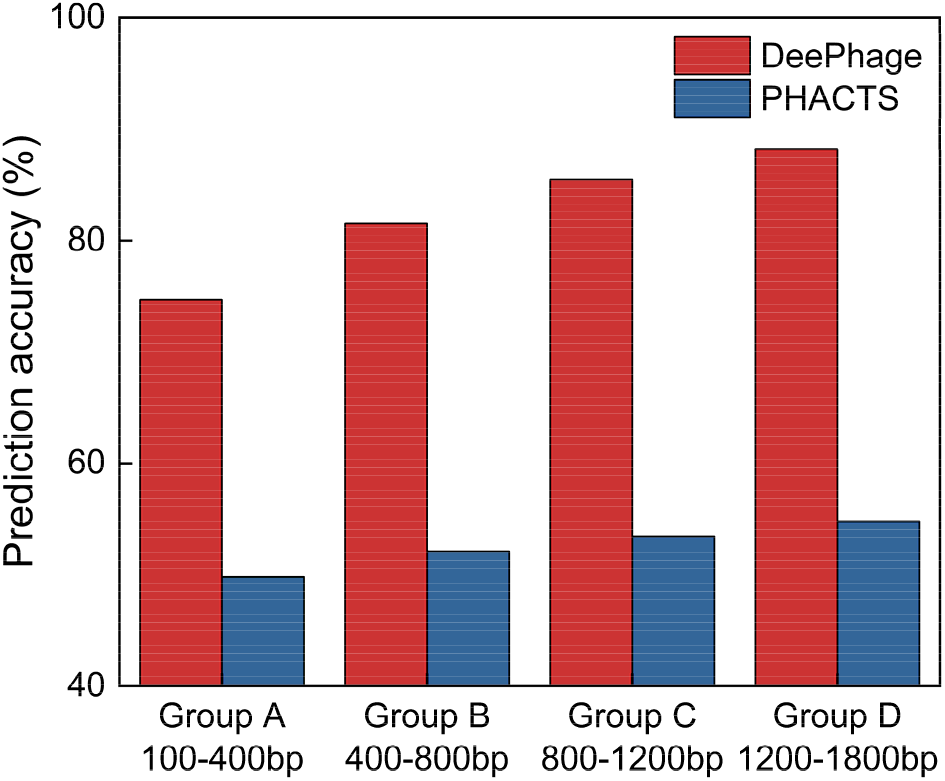
Comparison results of DeePhage and PHACTS in each length group.

In addition, we have evaluated the performances of DeePhage and PHACTS on the coding sequences (CDSs) from all 225 phage genomes. Since PHACTS could only process protein sequences, we extracted all CDSs from the genomes according to the GenBank annotation and each CDS was independently inputted to PHACTS (in the form of amino acid sequences) and DeePhage (in the form of nucleic acid sequences). We found that PHACTS can only achieve the *Acc* of 54.3%, which is also similar to random judgment results, while DeePhage achieves the *Acc* of 85.0%, more than 30% higher than that of PHACTS. Considering that the number of CDSs in each metagenomic fragment is very limited, PHACTS has actually a very limited ability to analyse metagenomic data especially when the complete genomes could not be reconstructed using these fragments. Overall, as the first tool designed for phage lifestyle classification from metagenomic data, DeePhage, a de novo tool using the deep learning algorithm, presents efficient prediction.

Also, DeePhage can handle large scale high-throughput data within an acceptable running time. In order to test, we recorded the runtime of DeePhage and PHACTS to predict 100 DNA sequences (converted to protein sequences for PHACTS) ranging from 100-1800 bp. DeePhage spends nearly 10 seconds 810 times faster than PHACTS using nearly 135 minutes, when tested on a virtual machine with the following configuration: CPU: Intel Core i7 4790; and Memory: 8G, DDR3. As for PHACTS, every sequence needs to be aligned and every prediction needs to be replicated ten times, while DeePhage could directly predict every sequence without any alignments. Therefore, DeePhage is much faster than PHACTS.

### Evaluation of DeePhage and PHACTS using real metavirome data

Although it was difficult to make exact evaluations using real data, some functional genes could help us to make an approximately effective assessment of our model. In this subsection, we used DeePhage to predict all the sequences in a metavirome data of bovine rumen [24] with 118918 contigs assembled by SPAdes (v3.13.0) [26]. As a result, 45.1% (53625/118918) of the contigs were predicted as virulent phage-derived contigs and 54.9% (65293/118918) as temperate phage-derived contigs. For assessment of the DeePhage’s prediction, we then collected the RefSeq viral protein database [35] as a reference. Since the viral proteins labelled of ‘excision’, ‘integration’, or ‘lysogeny’ are more likely to exist in temperate phages [14], we used those proteins to build an MTPD (mini temperate phage-derived) set containing 107 protein sequences. We then searched all 118918 contigs against the MTPD data using Blastx v2.7.1[36] and obtained 16 targeted contigs having homologous regions (e-value ≤ 1e-10, hits length ≥ 400). Those presented an extremely small proportion (16/118918), which confirms that there are a huge number of data having no reliable homologous regions of known databases. When it comes to DeePhage, 13 of 16 targeted contigs can be identified as temperate phage-derived contigs, while only 10 contigs can be classified as temperate phage-like contigs by PHACTS. It shows DeePhage performs better than PHACTS and has rather the potential to analyze newly sequenced phage data. However, the prediction scores being nearly 0.5 shows PHACTS actually made randomly inferring, while DeePhage having a majority of reliable scores and making better predictions. The information of 16 targeted contigs and predicted results by DeePhage and PHACTS was listed in Table 2.

**Table 2.** Information of 16 targeted contigs and predicted results by DeePhage and PHACTS. ‘Contig ID’ refers to the ID of 16 targeted contigs. ‘Identity’, ‘E-value’, and ‘Hits length’ refer to the alignment results using Blastx.

| Contig ID | Contig length (bp) | Identity (%) | E-value | Hits length | DeePhage |  | PHACTS |  |
| --- | --- | --- | --- | --- | --- | --- | --- | --- |
|  |  |  |  |  | prediction | score | prediction | score |
| 4 | 28516 | 26.32 | 1e-10 | 513 | temperate | 0.4308 | temperate | 0.4835 |
| 12 | 11212 | 27.23 | 2e-14 | 606 | temperate | 0.2022 | temperate | 0.4667 |
| 52 | 5349 | 29.89 | 1e-30 | 798 | temperate | 0.0127 | temperate | 0.4995 |
| 88 | 3734 | 24.68 | 1e-11 | 828 | temperate | 0.1326 | temperate | 0.4925 |
| 173 | 2530 | 26.22 | 2e-25 | 1044 | temperate | 0.0232 | virulent | 0.5000 |
| 223 | 2233 | 23.27 | 1e-12 | 834 | virulent | 0.6742 | virulent | 0.5161 |
| 1257 | 1029 | 29.87 | 1e-16 | 462 | virulent | 0.6044 | virulent | 0.5082 |
| 1639 | 921 | 23.96 | 2e-11 | 849 | temperate | 0.1589 | temperate | 0.4735 |
| 3299 | 702 | 28.18 | 8e-25 | 609 | temperate | 0.0303 | temperate | 0.4744 |
| 3326 | 699 | 30.88 | 9e-15 | 405 | temperate | 0.3095 | virulent | 0.5055 |
| 6405 | 549 | 25.14 | 1e-13 | 519 | temperate | 0.0222 | virulent | 0.5080 |
| 7704 | 514 | 39.86 | 2e-22 | 429 | temperate | 0.1451 | virulent | 0.5110 |
| 8130 | 503 | 36.05 | 3e-22 | 441 | temperate | 0.0512 | temperate | 0.4944 |
| 9804 | 470 | 31.69 | 2e-16 | 423 | temperate | 0.1596 | temperate | 0.4952 |
| 10819 | 454 | 38.61 | 2e-30 | 450 | temperate | 0.0574 | temperate | 0.4951 |
| 12636 | 430 | 34.04 | 2e-21 | 417 | virulent | 0.9784 | temperate | 0.4743 |

Further, we found that 16 contigs contain homologous regions of the functional proteins with the *e*-value lower than 1e-10, but they do not have high identity scores to these proteins (identity<50%). These results indicate that the 16 contigs are not close to the viral proteins from the database in the genetic relationship. These also show that the diversity of phages in the environmental samples might be much higher than that in the current database, and DeePhage can handle these novel phages. In fact, when we looked over the RefSeq viral protein database, we found that a large number of proteins are labelled as putative or hypothetical and the percentage of such proteins might be much higher than that of bacteria, which further demonstrates the diversity of phages.

Not only the several above-mentioned contigs but also the whole sequences could be taken into consideration in a full scope of the predictive ability of DeePhage. Using the whole genomes of 77 virulent and 148 temperate phages in our datasets, we annotated all the sequences in the virome data of bovine rumen by Blastn v2.7.1[36]. When setting the default parameters (the default e-value is 10), 118564 contigs could be annotated as virulent or temperate phage genomes by BLAST. Among those contigs, DeePhage distinguished 49.32% virulent and 57.69% temperate phage contigs with an average proportion of 54.4%. In comparison, PHACTS made an apparent preference for virulent contigs (68.2%) and for temperate contigs (28.9%). Although the proportion of virulent contigs was higher, PHACTS only received an average of 44.5%, which was much lower than DeePhage. Estimated on the level of entire real data, the superiority of DeePhage is certainly considerable.

To sum up, the evaluation of DeePhage using real metavirome data demonstrated DeePhage made much better and reliable predictions than PHACTS. As an ab initial tool, it may be concluded that DeePhage has a good ability to adapt to this diversity and has the potential to analyse newly sequenced phage data.

### An application of a cross-sectional study indicating that phage transformations impacting the change of gut microbiota structure

Viruses especially the phages contribute importantly to the gut microbiota structure. Particularly, temperate phages could be free from the genome of their bacterial hosts and then kill them driven by a suitable environment condition. While virulent phages directly attack their host. Therefore, such phage transformations would change the gut microbiota composition profile and community structure. However, it is hard to analyse this result entirely using the databased method because of the limitation of database and marker genes like 16S RNA. As a result, there are not effectively computational related tools. For example, alignment phage sequences to the known phage database using the traditional Blast program could just output some known phages without any new phages. Indeed, the number of unknown phages were extraordinarily huge. Fortunately, DeePhage now could detect phage transformations over the whole genomes of phages from the complete virome data. The downstream findings based on DeePhage could give us instructive insights into the function of phages in the gut microbiota.

In this subsection, we then designed a new strategy about how to use DeePhage to estimate the transformations of phages in the cross-sectional study. Specially, we analyse the virome data from ulcerative colitis (UC) patients and healthy people as an example to find out associations between phages and gut microbiota after having the disease. For phages in a community, owing to lack of marker genes like 16S RNA to detect their abundance or diversity, it is difficult to determine the association between the transformation of phages and the change of gut microbiota structure. Herein we collected 21 untargeted metagenomic samples (randomly selected) of UC patient guts and 21 (randomly selected) untargeted metagenomic samples of healthy human guts by Nielsen et al. [37]. In addition, we collected 54 virome samples (viral particles were enriched before sequencing) of UC patient guts (being diagnosed as a specific state) and 23 virome samples of healthy control by Norman et al. [38]. The accessions were provided in Additional File 1 (Table S1 and S2). We used SPAdes to assemble raw reads of each sample.

For each untargeted metagenomic sample, we first used PPR-Meta [10] to identify all the phage-derived contigs. The average percentage of phage contigs in metagenomic data of UC patient and healthy individual guts were similar (23.7% in UC patient and 25.7% in healthy human guts) without significant difference (*p*-value=0.170, the difference in location=0.021 and 95% confidence interval = (−0.007, 0.045) for two-sided Wilcoxon Rank-Sum test). For convenience, in the following text, phage contigs in gut microbiota annotated by PPR-Meta were referred to as computational phages while contigs from virome data were referred to as experimental phages. It was worth noting that experimental phages only included virulent phages and temperate phages in the lytic cycle. However, temperate phages in the lysogenic cycle could not be included, because temperate phages in the lysogenic cycle would integrate their genomes into host cells and would not assemble the viral particles. In contrast, computational phages included all kinds of phages.

We then used DeePhage to predict the lifestyle of the experimental phages. An average of 64.2% of the contigs were predicted as temperate phages in UC patients while 51.5% in healthy individuals with significant difference (*p*-value=0.001, difference in location=0.123 and 95% confidence interval = (0.054, 0.200) for two-sided Wilcoxon Rank-Sum test). This indicates that the proportion of temperate phages in UC patients’ gut was higher than in healthy individuals. However, we still could not infer the detailed transformations from this result, because both the decreased diversity of virulent phages and increased diversity of temperate phages in UC patients will lead to a higher proportion of temperate phages. More importantly, even if the number of virulent phages and temperate phages were the same in healthy individuals and UC patients, the proportion of temperate phages in experimental phages could also increase when more temperate phages were undergoing the transformation from the lysogenic cycle to the lytic cycle, in which they would assemble free viral particles. To make the population dynamics clearer, we further used DeePhage to predict the lifestyle of the computational phages. Surprisingly, an average of 57.5% and 56.9% of the contigs were predicted as temperate phages in UC patients and healthy individuals without significant difference (*p*-value=0.811, the difference in location=-0.003 and 95% confidence interval = (−0.036, 0.025) for two-sided Wilcoxon Rank-Sum test), indicating that the proportion of virulent phages and temperate phages in UC patients and healthy individuals were similar. Considering the results from computational phages and experimental phages together, it seemed that the higher proportion of temperate phages in experimental phages of UC patients might result from the part of temperate phages undergoing a transformation from the lysogenic cycle to the lytic cycle.

From these preliminary results, we inferred that the phage populations in UC patients were undergoing a kind of change that influence the gut microbiota structure, in which some kinds of temperate phages were transforming from prophages to free viral particles. To investigate the transforming temperate phages, we picked out all the temperate contigs annotated by DeePhage from the UC and healthy samples. Using all the phage genomes [39] as the database of the BLAST method (e-value ≤ 1e-10), 342 species of phages were existing in both Healthy and UC samples, and just 154 species, 99% of which were from the *Caudovirales* order, only existing in Healthy samples. As a comparison, we found out different phage contigs coming from 551 species that only existing in UC samples, which probably means there were more kinds of temperate phages in UC samples than in Healthy samples. Those phages could be classified into eleven families: *Siphoviridae, Herelleviridae, Podoviridae, Myoviridae, Ackermannviridae, Autographiviridae, Drexlerviridae, Tristromaviridae, Inoviridae, Microviridae, Sphaerolipoviridae.* The first seven families belong to the *Caudovirales* order, which accounts for nearly 97% (532/551) different species. Besides, a very small part (nine different species) is coming from *Microviridae* family. *Caudovirales* order and *Microviridae* family are dominated in human gut virome [40], meanwhile, they are more abundant in UC patients compared with household members and controls [41]. Especially, Norman et al. observed an increase in the richness of some members of the *Caudovirales* in UC patients [38]. Those supported our inference to a certain degree. The last several families lacking researchers’ concerns in the human gut could roughly be ignored. Since the release of prophages is often associated with the death of bacterial hosts, the activation of the temperate phages may be associated with the change of species composition. We can infer that after being illness more kinds of temperate *Caudovirales* phages turn into a lytic cycle and become free viral particles from the bacterial genomes, in consequence, such switch change the struct of microbiota by killing the bacterial host. Consistently, previous research showed that the species compositions of the bacteria community in UC patients were different from that of healthy individuals [42] and the virulent phages from the healthy core could be substituted by temperate phages [43]. All those discoveries indicated that maybe it was the temperate *Caudovirales* phages have a primary impact on human UC disease, which was also verified by us. However, DeePhage could not only detect well-studied phages, such as *Caudovirales* phages, but it also can trace any known and unknown phages to distinguish their lifestyles. With integrated data, we have access to disease conditions deeply.

To sum up, such a strategy being independent of databases may further provide insights into the specific and integral interactions between phages and bacterial hosts according to phage lifestyles, which could not have been found out before. Researchers can gain more valuable information about the disease process and facilitate the study of human disease.

## DISCUSSION

In this paper, we presented DeePhage as an effective tool to distinguish virulent phage-derived and temperate phage-derived sequences in metavirome data. Coding a DNA sequence, DeePhage needs no previously extracted features but use each nucleotide as input. There are some advantages. DeePhage can bypass using the information of some functional genes to make the judgment and directly and rapidly identify each DNA fragment being independent of assembling. Such a function is important because many novel phage genomes are difficult to reconstruct and the amount of sequences is large when focused on metagenomic data. CNN models here occupied the core strength of DeePhage for their excellent ability on feature extraction, which is hard to discover by statistics. As we can see, DeePhage gradually separates virulent and temperate phage-derived sequences along with deeper neural networks. DeePhage’s ability to distinguish two kinds of sequences is superior to PHACTS on the assessment of simulated data and real data. To be specific, DeePhage presents a huge improvement in prediction accuracy (nearly 30% higher on simulated data) and computational efficiency (almost 810 times faster). More importantly, DeePhage sheds new light on the phage transformations by tracing the variation of a specific type of phage. As we can see, the previous study speculated the possibility that the expansion of the *Caudovirales* phages is related to the activation of prophages in UC patients [44]. Fortunately, now we can be more convinced that more temperate *Caudovirales* phages are turning into a lytic cycle. We believe that there will be an increasing number of new discoveries, just like the problem mentioned before, on account of DeePhage. Afterward, DeePhage ultimately reduces the dependency on culture-dependent methods and promotes human disease research.

It is also interesting to explore the biological mechanism that helps DeePhage distinguish fragments from these two kinds of phage using the sequence signature. In our opinion, this may because virulent phages and temperate phages face different evolutionary pressures and therefore contain different sequence signatures, such as k-mer frequencies as we showed in Figure 1. Genome amelioration often occurs on foreign DNA, such as phage or plasmid, in the host cell and foreign DNA will change its sequence signatures according to the host chromosome to help it exist stably in the host cell [45]. The similarity of sequence signatures between foreign DNA and bacterial chromosome is often used to predict the bacterial host of the foreign DNA [11,12,45]. Since temperate phages will spend more time in the host cell, they may adjust their sequence signatures toward host chromosomes. Related researches also show that temperate phages do contain more similar sequence signatures to their hosts than virulent phages [21,46]. Therefore, we considered that the difference of sequence signatures played an important role for DeePhage to identify these two kinds of phages. To further prove this conjecture, we collected 120 bacterial genomes from RefSeq database [47] (the accession numbers can be seen in Additional File1, Table S4) and then used MetaSim to extract artificial contigs between 100 to 1800 bp. We observed how DeePhage would judge these bacterial sequences. Although the training set of DeePhage did not contain any bacterial sequences, DeePhage identified 79.3%, 84.7%, 84.9%, and 86.0% of the bacterial sequences as temperate phages in Group A, B, C, and D, respectively (the sequence length in each group was corresponding to Table 1). We considered that the reason why more than half of the bacterial sequences were identified as temperate phages was that bacteria contained similar sequence signatures with temperate phages. This phenomenon also demonstrates that using the information of sequence signatures may be the working principle of DeePhage.

DeePhage also has some limitations. Although prokaryotic viruses are dominant in virome samples, a few eukaryotic viruses could also be included. However, DeePhage cannot identify these sequences before distinguishing the lifestyle of each contig. Fortunately, the related tool that helps to distinguish prokaryotic and eukaryotic viruses has been developed recently [48] and we are also considering constructing a preprocessing module for DeePhage to filter out the eukaryotic viruses so that DeePhage can generate more reliable results for the downstream analysis.

In conclusion, to the best of our knowledge, DeePhage is the first tool that can directly judge each fragment as a virulent phage-derived or temperate phage-derived sequence for virome data in a fast way. Therefore, it is expected that DeePhage will be a powerful tool for researchers who are interested in the function of phage populations and phage-host interactions.

## Availability of supporting data and materials

The artificial contigs, related scripts, and original results are available at http://cqb.pku.edu.cn/ZhuLab/DeePhage/data/. All the other data are available at corresponding references mentioned in the main text.

## Availability of supporting source code and requirements

Project name: DeePhage.

Project home page: http://cqb.pku.edu.cn/ZhuLab/DeePhage or https://github.com/shufangwu/DeePhage.

Operating system: The code of DeePhage was written on Linux. We optimized the program in a virtual machine; thus, DeePhage is platform independent.

Programming language: python, matlab

Other requirements: no other requirements are needed if running in the virtual machine. If not, Python 3.6.7, TensorFlow 1.4.0, Keras 2.1.3, numpy 1.16.4, h5py 2.9.0 and MATLAB Component Runtime 2018a (for free) are needed. MATLAB is not necessary.

License: GPL-3.0.

RRID: SCR_019243

## Additional files

Additional file 1: Figure S1. The architectures of six different models; Table S1. The Sn, Sp, and Acc of six different models; Figure S2. The ROC curves and AUC scores of DeePhage performances in each set of five-fold cross-validation; Table S2. The accession numbers of 21 untargeted metagenomic samples of the healthy human gut and 21 untargeted metagenomic samples of UC patients’ gut; Table S3. The accession numbers of 23 virome samples of the healthy human gut and 54 virome samples of UC patients’ gut; Table S4. The accession numbers of 120 bacterial genomes from RefSeq database.

## Authors’ contributions

H.Q.Z. and S.F.W. proposed and designed the study. J.T. constructed the datasets. S.F.W. and Z.C.F. optimized the code. M.L., C.H.W., and Q.G. contributed to the analysis. C.M.X and X.Q.J helped to test the results. S.F.W. and H.Q.Z. wrote and revised the manuscript, and all authors proofread and improved the manuscript.

## ACKNOWLEDGEMENT

We thank Dr. Li Qu, Luotong Wang, Man Zhou and Chuan He of Peking University for their helpful discussions. Part of the analysis was performed on the High Performance Computing Platform of the Center for Life Science of Peking University.

