## Additional File for "Distinguish virulent and temperate phage-derived sequences in metavirome data with a deep learning approach"

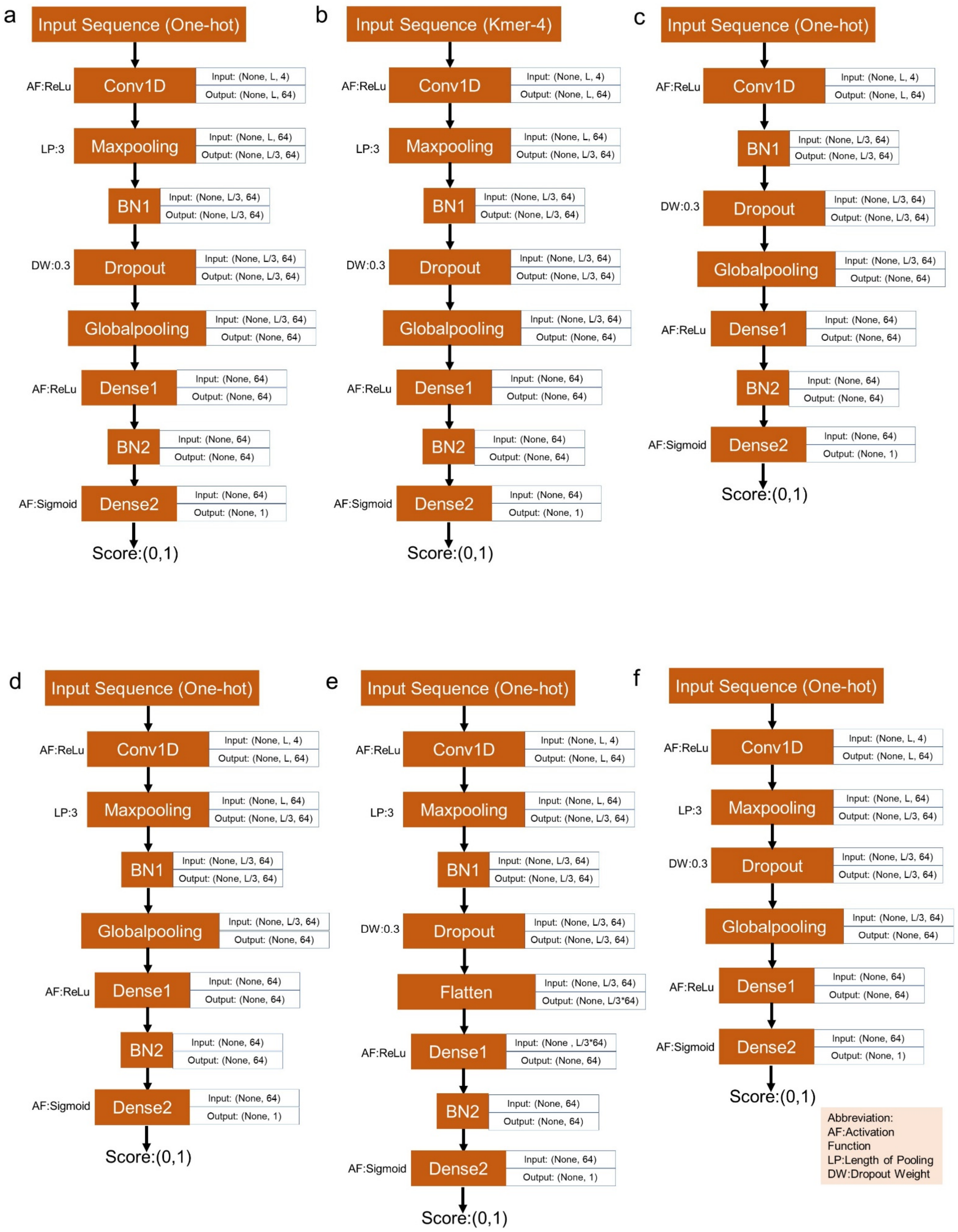
a
Input Sequence (One-hot)
AF:ReLu
Conv1D
Input: (None, L, 4)
Output: (None, L, 64)
LP:3
Maxpooling
Input: (None, L, 64)
Output: (None, L/3, 64)
BN1
Input: (None, L/3, 64)
Output: (None, L/3, 64)
DW:0.3
Dropout
Input: (None, L/3, 64)
Output: (None, L/3, 64)
Globalpooling
Input: (None, L/3, 64)
Output: (None, 64)
AF:ReLu
Dense1
Input: (None, 64)
Output: (None, 64)
BN2
Input: (None, 64)
Output: (None, 64)
AF:Sigmoid
Dense2
Input: (None, 64)
Output: (None, 1)
Score:(0,1)
b
Input Sequence (Kmer-4)
AF:ReLu
Conv1D
Input: (None, L, 4)
Output: (None, L, 64)
LP:3
Maxpooling
Input: (None, L, 64)
Output: (None, L/3, 64)
BN1
Input: (None, L/3, 64)
Output: (None, L/3, 64)
DW:0.3
Dropout
Input: (None, L/3, 64)
Output: (None, L/3, 64)
Globalpooling
Input: (None, L/3, 64)
Output: (None, 64)
AF:ReLu
Dense1
Input: (None, 64)
Output: (None, 64)
BN2
Input: (None, 64)
Output: (None, 64)
AF:Sigmoid
Dense2
Input: (None, 64)
Output: (None, 1)
Score:(0,1)
c
Input Sequence (One-hot)
AF:ReLu
Conv1D
Input: (None, L, 4)
Output: (None, L, 64)
BN1
Input: (None, L/3, 64)
Output: (None, L/3, 64)
DW:0.3
Dropout
Input: (None, L/3, 64)
Output: (None, L/3, 64)
Globalpooling
Input: (None, L/3, 64)
Output: (None, 64)
AF:ReLu
Dense1
Input: (None, 64)
Output: (None, 64)
BN2
Input: (None, 64)
Output: (None, 64)
AF:Sigmoid
Dense2
Input: (None, 64)
Output: (None, 1)
Score:(0,1)
d
Input Sequence (One-hot)
AF:ReLu
Conv1D
Input: (None, L, 4)
Output: (None, L, 64)
LP:3
Maxpooling
Input: (None, L, 64)
Output: (None, L/3, 64)
BN1
Input: (None, L/3, 64)
Output: (None, L/3, 64)
Globalpooling
Input: (None, L/3, 64)
Output: (None, 64)
AF:ReLu
Dense1
Input: (None, 64)
Output: (None, 64)
BN2
Input: (None, 64)
Output: (None, 64)
AF:Sigmoid
Dense2
Input: (None, 64)
Output: (None, 1)
Score:(0,1)
e
Input Sequence (One-hot)
AF:ReLu
Conv1D
Input: (None, L, 4)
Output: (None, L, 64)
LP:3
Maxpooling
Input: (None, L, 64)
Output: (None, L/3, 64)
BN1
Input: (None, L/3, 64)
Output: (None, L/3, 64)
DW:0.3
Dropout
Input: (None, L/3, 64)
Output: (None, L/3, 64)
Flatten
Input: (None, L/3, 64)
Output: (None, L/3*64)
AF:ReLu
Dense1
Input: (None , L/3*64)
Output: (None, 64)
BN2
Input: (None, 64)
Output: (None, 64)
AF:Sigmoid
Dense2
Input: (None, 64)
Output: (None, 1)
Score:(0,1)
f
Input Sequence (One-hot)
AF:ReLu
Conv1D
Input: (None, L, 4)
Output: (None, L, 64)
LP:3
Maxpooling
Input: (None, L, 64)
Output: (None, L/3, 64)
DW:0.3
Dropout
Input: (None, L/3, 64)
Output: (None, L/3, 64)
Globalpooling
Input: (None, L/3, 64)
Output: (None, 64)
AF:ReLu
Dense1
Input: (None, 64)
Output: (None, 64)
AF:Sigmoid
Dense2
Input: (None, 64)
Output: (None, 1)
Score:(0,1)
Abbreviation:
AF:Activation Function
LP:Length of Pooling
DW:Dropout Weight

**Figure S1.** The architectures of six different models. (a). The model architecture of DeePhage. (b). The model architecture of the Kmer-4 model. Based on DeePhage, Kmer-4 just used 4-mer frequencies to encode input sequences. (c). The model architecture of the No-Maxpooling model. Based on DeePhage, No-Maxpooling just took out of the Maxpooling layer. (d). The model architecture of the No-Dropout model. Based on DeePhage, No-Dropout just took out of the Dropout layer. (e). The model architecture of the No-Globalpooling model. Based on DeePhage, No-Globalpooling just took out of the Globalpooling layer and used a Flatten layer (flattens the input into one dimension) instead. (f). The model architecture of the No-BN model. Based on DeePhage, No-BN just took out of two BN layers.

**Table S1.** The Sn, Sp, and Acc of six different models. The length of the input sequences is 1800 and all the hyper-parameters of each model are nearly the same as DeePhage.

| Model | DeePhage | Kmer-4 | No-Maxpooling | No-Dropout | No-Globalpooling | No-BN |
|---|---|---|---|---|---|---|
| *Sn* (%) | 81.3 | 89.7 | 77.7 | 71.3 | 72.2 | 74.5 |
| *Sp* (%) | 92.6 | 27.8 | 94.6 | 96.5 | 82.8 | 89.7 |
| *Acc* (%) | 87.5 | 61.6 | 86.9 | 85.1 | 78.0 | 82.8 |

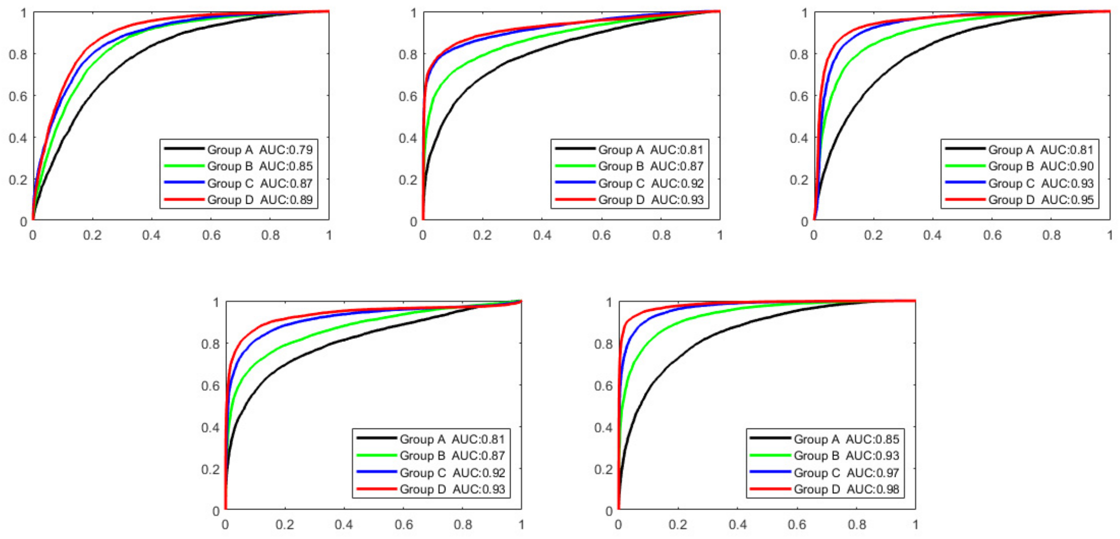


**Figure S2.** The ROC curves and AUC scores of DeePhage performances in each set of five-fold cross-validation.

**Table S2.** The accession numbers of 21 untargeted metagenomic samples of the healthy human gut and 21 untargeted metagenomic samples of UC patients' gut.

| Accessions of Health Control | Accessions of Ulcerative Colitis |
|---|---|
| ERS199004 | ERS199177 |
| ERS199005 | ERS199178 |
| ERS199007 | ERS199179 |
| ERS199010 | ERS199180 |
| ERS199012 | ERS199181 |
| ERS199016 | ERS199182 |
| ERS199018 | ERS199183 |
| ERS199021 | ERS199184 |
| ERS199023 | ERS199185 |
| ERS199026 | ERS199186 |
| ERS199027 | ERS199187 |
| ERS199029 | ERS199188 |
| ERS199033 | ERS199189 |
| ERS199035 | ERS199190 |
| ERS199036 | ERS199191 |
| ERS199037 | ERS199192 |
| ERS199038 | ERS199193 |
| ERS199039 | ERS199194 |
| ERS199040 | ERS199195 |
| ERS199041 | ERS199196 |
| ERS199042 | ERS199197 |

**Table S3.** The accession numbers of 23 virome samples of the healthy human gut and 54 virome samples of UC patients' gut.

| Accessions of Health Control | Accessions of Ulcerative Colitis |
|---|---|
| ERS698757 | ERS698653 |
| ERS698759 | ERS698654 |
| ERS698760 | ERS698656 |
| ERS698761 | ERS698657 |
| ERS698762 | ERS698659 |
| ERS698763 | ERS698660 |
| ERS698764 | ERS698661 |
| ERS698765 | ERS698662 |
| ERS698766 | ERS698664 |
| ERS698767 | ERS698665 |
| ERS698768 | ERS698666 |
| ERS698769 | ERS698667 |
| ERS698770 | ERS698668 |
| ERS698771 | ERS698669 |

| | |
|---|---|
| ERS698772 | ERS698670 |
| ERS698773 | ERS698671 |
| ERS698774 | ERS698672 |
| ERS698775 | ERS698673 |
| ERS698776 | ERS698674 |
| ERS698777 | ERS698682 |
| ERS698778 | ERS698686 |
| ERS698779 | ERS698687 |
| ERS698780 | ERS698689 |
| / | ERS698693 |
| / | ERS698694 |
| / | ERS698695 |
| / | ERS698696 |
| / | ERS698697 |
| / | ERS698699 |
| / | ERS698701 |
| / | ERS698706 |
| / | ERS698708 |
| / | ERS698709 |
| / | ERS698710 |
| / | ERS698713 |
| / | ERS698715 |
| / | ERS698716 |
| / | ERS698753 |
| / | ERS698754 |
| / | ERS698755 |
| / | ERS698756 |
| / | ERS698758 |
| / | ERS698798 |
| / | ERS698799 |
| / | ERS698800 |
| / | ERS698802 |
| / | ERS698803 |
| / | ERS698804 |
| / | ERS698806 |
| / | ERS698808 |
| / | ERS698811 |
| / | ERS698814 |
| / | ERS698815 |
| / | ERS698822 |

**Table S4.** The accession numbers of 120 bacterial genomes from RefSeq database.

| | | |
|---|---|---|
| NC_016603.1 | NC_011661.1 | NC_003112.2 |
| NC_008570.1 | NC_014121.1 | NC_005042.1 |
| NC_003063.2, NC_003062.2 | NC_004668.1 | NC_002516.2 |
| NC_006840.2, NC_006841.2 | NC_017960.1 | NC_002947.4 |
| NC_014318.1 | NC_011750.1 | NC_007005.1 |
| NC_000918.1 | NC_018658.1 | NC_004578.1 |
| NC_003997.3 | NC_002695.2 | NC_007494.2, NC_007493.2 |
| NC_005945.1 | NC_017634.1 | NC_005027.1 |
| NC_004722.1 | NC_000913.3 | NC_007643.1 |
| NC_000964.3 | NC_011751.1 | NC_000963.1 |
| NC_005957.1 | NC_009613.3 | NC_007677.1 |
| NC_006347.1 | NC_006570.2 | NC_003198.1 |
| NC_004663.1 | NC_003454.1 | NC_003197.2 |
| NC_014638.1 | NC_014644.1 | NC_004347.2 |
| NC_004307.2 | NC_002939.5 | NC_007606.1 |
| NC_019382.1 | NC_005125.1 | NC_004337.2 |
| NC_018828.1 | NC_000907.1 | NC_012587.1 |
| NC_002929.2 | NC_000915.1 | NC_009636.1 |
| NC_001318.1 | NC_017384.1 | NC_003047.1 |
| NC_013172.1 | NC_015663.1 | NC_007795.1 |
| NC_004463.1 | NC_016845.1 | NC_004461.1 |
| NC_002528.1 | NC_006814.3 | NC_004116.1 |
| NC_006349.2, NC_006348.1 | NC_008526.1 | NC_013853.1 |
| NC_006350.1, NC_006351.1 | NC_004567.2 | NC_004350.2 |
| NC_002163.1 | NC_007929.1 | NC_003098.1 |
| NC_002696.2 | NC_002662.1 | NC_002737.2 |
| NC_011916.1 | NC_002942.5 | NC_009009.1 |
| NC_000922.1 | NC_004343.2, NC_004342.2 | NC_012926.1 |
| NC_010287.1 | NC_003210.1 | NC_003888.3 |
| NC_000117.1 | NC_006055.1 | NC_013522.1 |
| NC_002932.3 | NC_014923.1 | NC_011296.1 |
| NC_010175.1 | NC_007644.1 | NC_004113.1 |
| NC_009089.1 | NC_002677.1 | NC_000853.1 |
| NC_003030.1 | NC_000962.3 | NC_006461.1 |
| NC_009495.1 | NC_002945.4 | NC_002967.9 |
| NC_009698.1 | NC_010397.1 | NC_002505.1, NC_002506.1 |
| NC_003450.3 | NC_008596.1 | NC_004605.1, NC_004603.1 |
| NC_002971.4 | NC_005364.2 | NC_003902.1 |
| NC_001263.1, NC_001264.1 | NC_000912.1 | NC_008800.1 |
| NC_002937.3 | NC_002946.2 | NC_003143.1 |
